# Differential genomic and transcriptomic events associated with high-grade transformation of Chronic Lymphocytic Leukemia

**DOI:** 10.1101/644542

**Authors:** Jenny Klintman, Basile Stamatopoulos, Katie Ridout, Toby A. Eyre, Laura Lopez Pascua, Niamh Appleby, Samantha J. L. Knight, Helene Dreau, Niko Popitsch, The HICF2 Consortium, Mats Ehinger, Jose I. Martín-Subero, Elias Campo, Robert Månsson, Davide Rossi, Jenny C. Taylor, Dimitrios V. Vavoulis, Anna Schuh

## Abstract

The transformation of chronic lymphocytic leukemia (CLL) to high-grade diffuse large B-cell lymphoma (DLBCL), also called Richter’s Syndrome (RS), is a rare cancer with dismal prognosis. Drug discovery for RS is hampered by the lack of suitable experimental models, and effective therapies remain elusive rendering RS an area of high unmet clinical need. We performed whole genome sequencing (WGS) to interrogate paired CLL and RS samples from 17 patients enrolled in a prospective multicenter Phase 2 clinical trial (CHOP-OR) and we found that subclones affected by mutations in MAPK and PI3K pathways show a high expansion probability during transformation. We also demonstrate for the first time that non-coding mutation clusters in a *PAX5* enhancer, situated 330kb upstream from the transcription initiation site, correlate with transformation. Finally, we confirm our findings by employing targeted DNA sequencing (TGS) and RNA expression profiling on an extended cohort of 38 patients.

**Statement of significance:** Through integrated analysis of WGS, TGS and RNA expression data, we identified drivers of transformation not previously implicated in RS, which can be targeted therapeutically and tested in the clinic. Our results have informed the design of a new clinical platform study, which is now open to recruitment in the UK.

## Introduction

The transformation of chronic lymphocytic leukemia (CLL) into an aggressive, high-grade diffuse large B-cell lymphoma (DLBCL) called Richter’s syndrome (RS) occurs in 2-15% of CLL patients^1–4^. Based on the annual prevalence of CLL in the UK (5220 cases)^5,6^, RS is a rare cancer with a prevalence of approximately 100 cases per year (0.17 per 100,000), while its incidence increases significantly in heavily pre-treated patients^7^. RS has an overall survival from diagnosis of 5.9 to 11.4 months using standard-of-care therapy with cyclophosphamide, doxorubicin, vincristine, prednisolone and rituximab (CHOP-R) and therefore remains an area of high unmet clinical need^3,5,8^. Although small molecule inhibitors (SMIs) targeting the B-cell receptor (BCR) pathway and BCL2^9–11^ have significantly improved the outlook of patients with CLL, RS is a frequent cause of SMI failure^4,11–14^. Although RS shares the histological characteristics of DLBCL, it has a distinct molecular profile compared to *de novo* DLBCL as 90% of cases described in the literature carry molecular lesions that emerge from a CLL-related clone in at least one of the following genes: *TP53 (60-80%), CDKN2A (30%), MYC (30%), MGA (7%)* or *NOTCH1 (30%)*^3,15–19^.

Drug discovery for RS is hampered by the lack of suitable *in vitro* or *in vivo* models. Novel agents and their combinations (including the checkpoint PD-1 inhibitor pembrolizumab, second generation BCR inhibitors acalabrutinib and umbralisib, and the XPO-1 inhibitor Selinexor^3,8,20^) are currently under investigation. We previously conducted the first and so far only prospective multi-center Phase II study (CHOP-OR) exclusively for patients with treatment-naïve, biopsy-confirmed DLBCL-type RS^21^. We concluded that the TP53-independent properties of the Type 2 anti-CD20 antibody ofatumumab in combination with CHOP followed by ofatumumab maintenance therapy did not improve patient outcomes compared to historical controls. The study protocol included extensive sample collection aimed at identifying clinically-testable putative transformation drivers.

In this study, we present results from the first integrative WGS analysis of paired CLL and RS samples using a combination of coding and non-coding single nucleotide variants (SNVs), small insertions or deletions (InDels) and copy number aberrations (CNAs), as well as TGS and RNA expression profiling to interrogate genes and pathways involved in the transformation to RS. Using a combined cohort of 55 RS patients (the second largest so far), we managed to identify multiple targetable pathways as putative transformation drivers for further clinical evaluation.

## Results

### Detection of genetic variants in the CHOP-OR discovery cohort

#### Mutation Burden

Paired germline, CLL and lymphoma samples from 17 subjects in the CHOP-OR study^21^ underwent WGS. The clinical characteristics of the subjects and results from IgHV analysis on CLL and RS tumor samples are presented in Supplementary Table 1.

Despite optimization of DNA extraction methods, formalin-fixation induces DNA artefacts that increase the false positive mutation rate^22^. We therefore applied a stringent variant allele frequency (VAF) filter (see Methods) on all FFPE-derived data, and chose a genomic region unlikely to be affected by RS driver mutations (i.e. the non-rearranged T-cell receptor-{C,V,J} gene locus) as an internal control for effective filtering. As expected, no somatic variants were detected in CLL or RS samples across this locus after filtering (Supplementary Figure 1).

In total, 884 somatic non-synonymous variants remained in CLL and RS samples (Figure 2A and B; Supplementary Tables 2 and 3). The mean read depths over the alternate alleles were 82X (±30) and 76X (±50) in the CLL samples (n=354 total variants) and RS samples (n=530 total variants), respectively. The mean VAFs for the detected variants were 0.27 (±0.16; range 0.05-0.89) and 0.32 (±0.16; range 0.10-0.88) in the CLL and RS samples, respectively.

**Figure 1:**
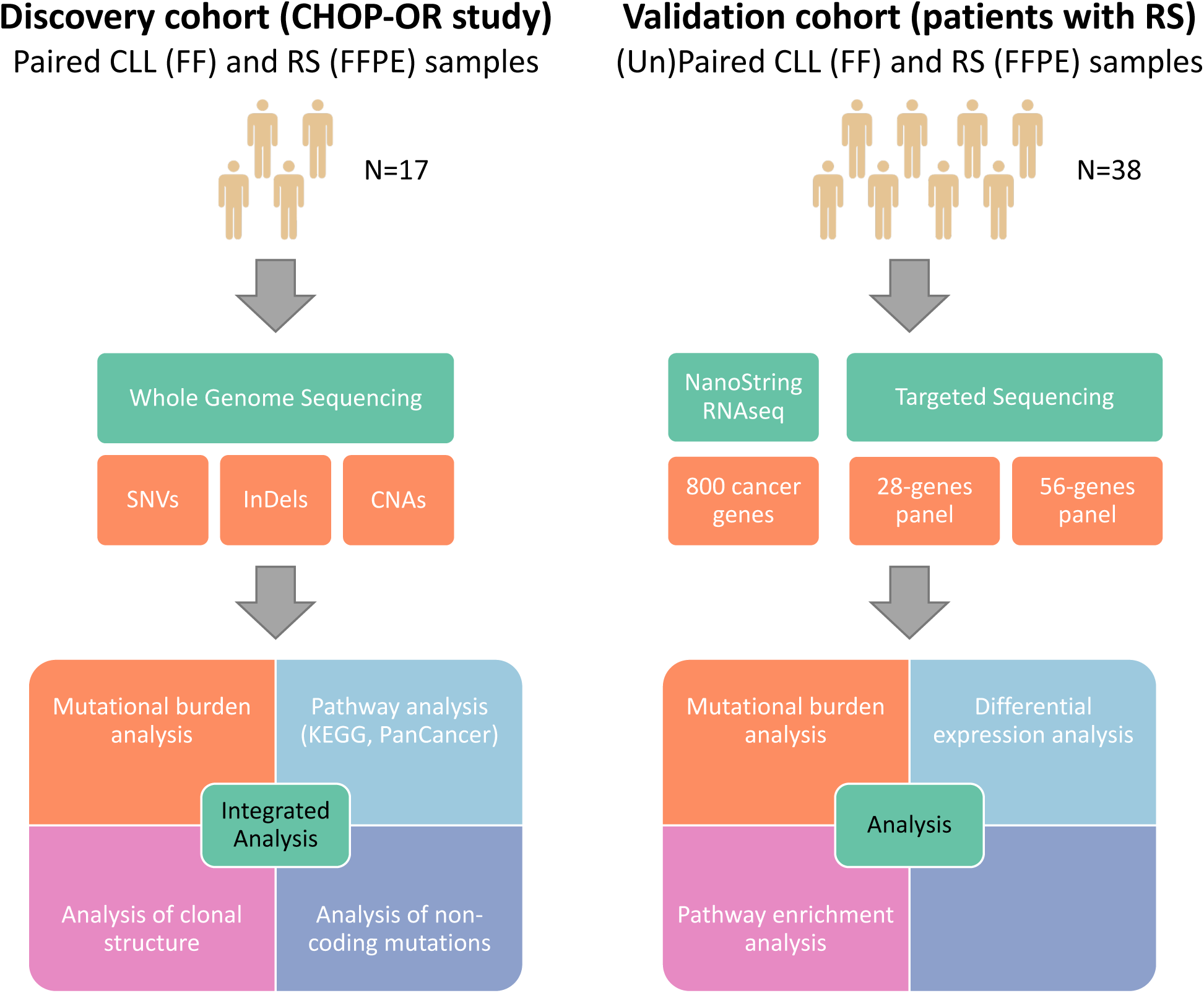
Overview of the study. SNVs, InDels and CNVs were identified in pairs of CLL and lymph-node samples from 17 CHOP-OR patients^21^ with Richter’s syndrome using whole genome sequencing. Integrated analysis of mutational burden and clonal structure based on these data was followed by differential gene expression and enrichment analysis and mutational burden analysis based on targeted sequencing on independent cohorts of 38 subjects with Richter’s syndrome.

**Figure 2:**
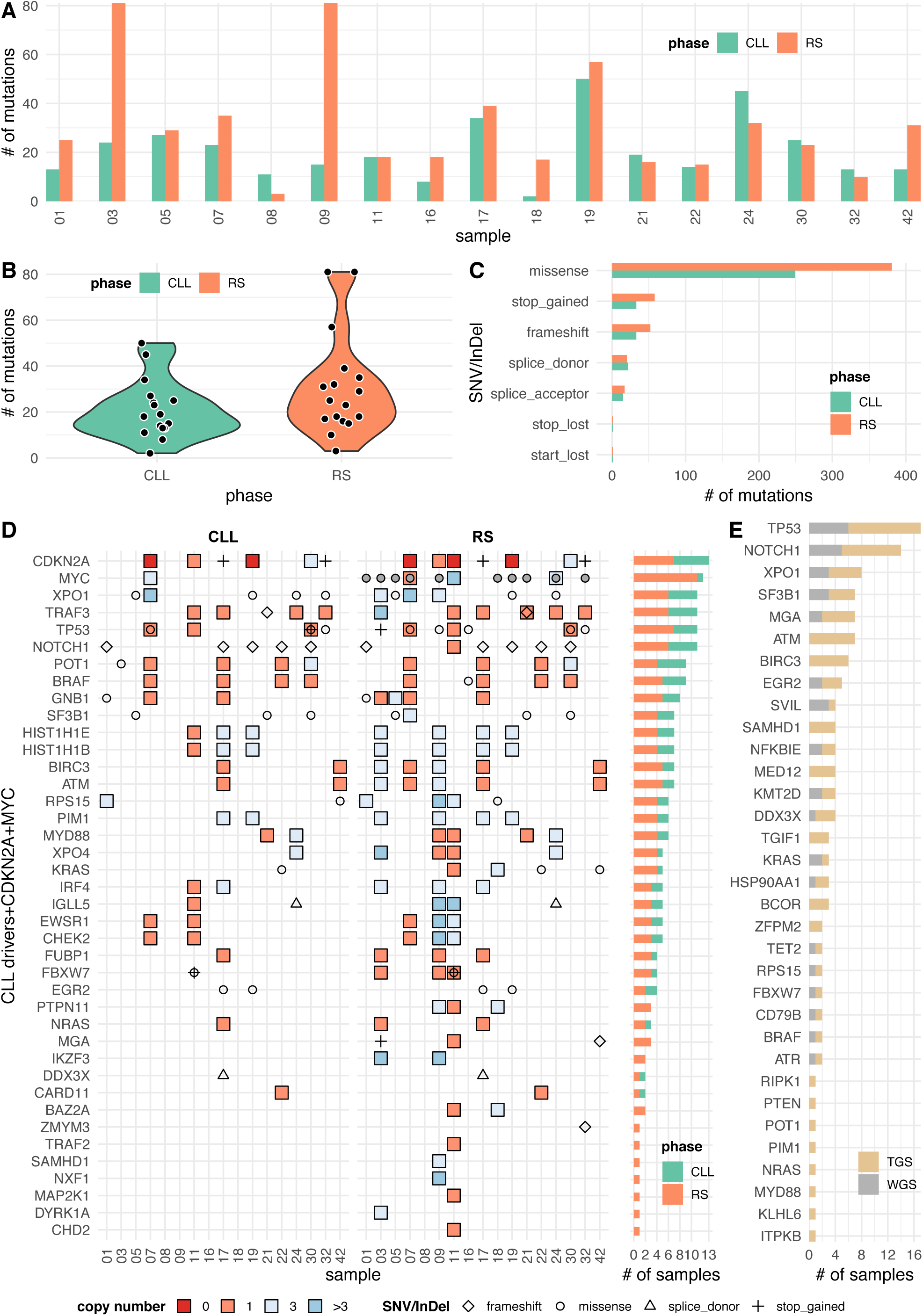
SNVs, InDels and CNVs in the cohort of 17 patients with Richter’s syndrome. A) Number of SNVs and InDels in the CLL versus the RS phase in each of the 17 patients. B) Number of mutations in the CLL vs RS phase across all samples (P=0.031; one-sided Wilcoxon signed rank test with continuity correction). C) Consequences of filtered SNVs and InDels and their distribution between the CLL and RS phases. D) Distribution of SNVs, InDels and CNVs across samples and phases in CLL driver genes, CDKN2A and MYC. 44 CLL driver genes were considered^23^, only 38 of which harbored an aberration in our cohort. Grey dots indicate cases with *MYC* over-expression in the RS phase (range: 33.9% in case 18 to 754.9% in case 07; mean: 147.8%). *MYC* deregulation (i.e. over-expression) was confirmed using differential RNA expression analysis. E) Frequency of mutated genes identified using targeted sequencing (n=38) merged with results from WGS analysis (n=17). Only SNVs and InDels are illustrated.

A significantly larger SNVs and InDels burden was detected in RS (31.2±22.5 variants per subject) compared to CLL phase (20.8±12.7 variants per subject) (Figure 2B; *p*=0.031; one-sided Wilcoxon signed rank test with continuity correction), and significantly more genes were mutated in RS (n=457 mutated genes; 29.6±21.3 mutated genes per subject) compared to CLL (n=306 mutated genes; 19.8±12.4 mutated genes per subject) (*p*=0.022; same test as above). Most mutated genes and variants detected were non-recurrent (i.e. only detected once and in one subject). Missense mutations were most common. No specific type of mutation was associated with disease phase (Figure 2C). The same pattern was found across all coding mutations and most patients, although three patients (03, 09 and 19) had a particularly high mutational burden.

Several types of acquired copy number aberrations were detected in both CLL and RS phases, including homozygous (CN=0) and heterozygous (CN=1) copy number loss, copy number gain (CN=3) and high copy number gain (CN>3) with or without loss of heterozygosity (LOH) (Supplementary Figure 2 and Supplementary Table 4). The number of genes affected by a CNA varied largely between disease phases and subjects. In some cases (e.g. no’s 03, 09 and 11), the total number of genes affected in the RS phase was close to or higher than 20K genes (Supplementary Figure 2A). Heterozygous copy number loss was the most common CNA, followed by CNA gain (Supplementary Figure 2B).

#### Recurrently mutated genes in the RS phase

Next, we performed integrated analysis of all acquired SNVs/InDels and CNAs in 44 CLL drivers^23^ (Figure 2D; Supplementary Table 5). Consistent with previous studies^3,15,16,24^, we found lesions in the *TP53, NOTCH1* and *CDKN2A* genes to approximately the same extent as in comparable RS cohorts (Figure 2D; Supplementary Tables 2 and 4). In our cohort, at least one of these genomic lesions was found in 9 CLL samples (52.9%) compared to 13 RS samples (76.5%).

Overall, we found mutations in 38 CLL driver genes (38/44; 86.4%), either in the RS phase only, or in both phases. Thirteen CLL samples (76.5%) and all but one RS samples (n=16; 94.1%) had at least one CLL driver mutation (Figure 2D and E). As previously reported for RS, the *NOTCH1* (n=6; 35.3%) and *TP53* (n=7; 41.2%) genes frequently harboured genetic lesions (Figure 2D). In addition, we found the *XPO1* (n=6; 35.3%) and *TRAF3* (n=6; 35.3%) genes to be mutated at a similar frequency in the RS phase (Figure 2D). Mutations in four CLL driver genes were recurrently detected in the RS phase only: *PTPN11* (n=3; 17.6%), *MGA* (n=3; 17.6%), *IKZF3* (n=2; 11.8%) and *BAZ2A* (n=2; 11.8%) (Figure 2D).

Furthermore, because RS is often associated with primary chemotherapy resistance, we investigated mutations in a custom set of 21 DNA damage response (DDR) genes (Supplementary Table 5; Supplementary Figure 3) and we found 13 out of 17 RS samples (76.5%) compared to 9 out of 17 CLL samples (52.9%) with mutations in at least one DDR gene. Of the recurrently mutated DDR genes, *BRIP1, RAD51C* and *RAD51D* acquired lesions in the transition from CLL to RS, but overall, DDR genes were found to be mutated in both disease phases (Supplementary Figure 3). Notice that 4 out of the 21 DDR genes (*RAD50, FANCA, ERCC3* and *MRE11*) were not found to harbour any lesions in either CLL or RS and they were omitted from Supplementary Figure 3.

#### Genes with mutations exclusively found or clonally expanded in the RS phase

In order to identify genes and specific variants associated with malignant transformation from CLL to RS, we next focused the analysis on recurrent SNVs/InDels that either occurred exclusively in RS or that clonally expanded in RS compared to the CLL phase of the same patient. We identified 33 recurrently mutated genes in RS harboring 80 unique SNVs/InDels (Supplementary Table 6). Among these, 51 variants (63.8%) across 30 genes (90.9%) were exclusively present in RS or showed clonal expansion during transformation to RS. Seventeen of these 30 genes (56.7%; indicated in red) were found recurrently mutated: *ABCD1P3, CSMD3, DND1, DNER, DST, DUSP2, IGSF3, IRF2BP2, KMT2C, MGA, PRAMEF1, RHPN2, SLC9B1, SVIL, TP53, VEZT, WWP1*. Most variants detected in this group of genes were non-recurrent in our cohort (Supplementary Table 6), but 7 among them (13.7%; indicated in red) were found to recur in two samples each: chr1:g.117156459C>T (in gene *IGSF3*), chr10:g.29784072G>C (in gene *SVIL*), chr16:g.32487123T>C (in gene *ABCD1P3*), chr19:g.33490566T>C (in gene *RHPN2*), chr4:g.103826757T>C and chr4:g.103826769G>A (in gene *SLC9B1*), and chr5:g.140050940C>T (in gene *DND1*). The only other recurrent mutations in RS were chr9:g.139390648CAG>C (in *NOTCH1*; n=5 samples; 29.4%) and chr2:g.61719472C>T (in *XPO1*; n=3 samples; 17.6%), both of which are well-known hotspot mutations.

#### Targeted sequencing on the extended cohort

In order to extend the analysis of gene mutation recurrence to a larger cohort, we performed targeted sequencing on 38 additional cases. After combining SNVs/InDels detected in the discovery and in the independent cohort (n=55), *TP53, NOTCH1, XPO1, ATM, SF3B1, MGA, BIRC3* and *EGR2, all associated with poor risk CLL*, were the most commonly mutated genes (Figure 2E; Supplementary Table 7). In four genes, specific variants were recurrent: *NOTCH1* chr9:g.139390648CAG>C, *EGR2* chr10:g.64573248G>T, *XPO1* chr2:g.61719472C>T and *TP53* chr17:g.7578212G>A (Supplementary Tables 2 and 8). The *EGR2* variant has never been reported in RS before, but other *EGR2* mutations have^25^. The *TP53* chr17:g.7578212G>A stop variant has never been reported in any hematological malignancies before, but it has been reported in more than 80 cases of carcinoma^26^.

### Pathway analysis

Up to this point, we have examined the overall mutation burden of RS samples compared to CLL samples with respect to all different types of genomic aberrations, but the combined impact of these aberrations on cellular pathways is unknown. We therefore expanded our analysis to include extensive gene lists covering previously identified CLL drivers, DNA damage response (DDR) genes, cell signaling, cell cycle, apoptosis and general cancer pathways (Supplementary Table 5). In total, we examined 45 different gene sets for enrichment of genomic aberrations (somatic SNVs/InDels and CNAs) in the RS and CLL phases. Pathways more likely to harbor both a somatic SNV/InDel and a CNA in the RS phase (when compared to CLL) with a probability higher than 95% (false discovery rate or FDR<5%; see Supplementary Statistical Methods) were considered differentially mutated (Figure 3A). Using this approach, we found that the MAPK pathway had a higher burden (FDR=3.12%) in RS compared to CLL (Figure 3A). Lesions in at least one MAPK pathway gene were detected in 9 (52.9%) CLL and 15 (88.2%) RS samples (Figure 3B). In addition, the phosphoinositide 3-kinase (PI3K) pathway fell just below the 95% cut-off (FDR=5.05%; Figure 3A and Supplementary Figure 4). At least one PI3K pathway gene lesion was detected in 10 (58.8%) CLL and 14 (82.4%) RS samples. Furthermore, the analysis confirmed that neither CLL drivers nor DDR genes were more often mutated in RS compared to CLL (Figure 3A). Lesions (SNVs, InDels and/or CNAs) were detected in 14 of 270 (5.2%) MAPK genes tested. In the majority of these (n=13; 92.9%), mutations were detected in at least two separate RS samples (recurrently mutated genes; Figure 3B). *TP53* (n=7; 41.2%) and *BRAF* (n=5; 29.4%) were most frequently mutated in RS, followed by *DUSP5, RAF1, MAP2K2, KRAS, CACNA1D* with four (23.5%) mutated samples each. Among these genes, *KRAS*^25^, *BRAF*^27^ and *TP53*^15,16,24^ have been previously reported mutated in other RS cohorts. Furthermore, similar to the CLL8 trial^28^, our analysis could not confirm association between the NOTCH signaling pathway and transformation from CLL to RS^16^ (FDR=51.23%; Figure 3A).

**Figure 3:**
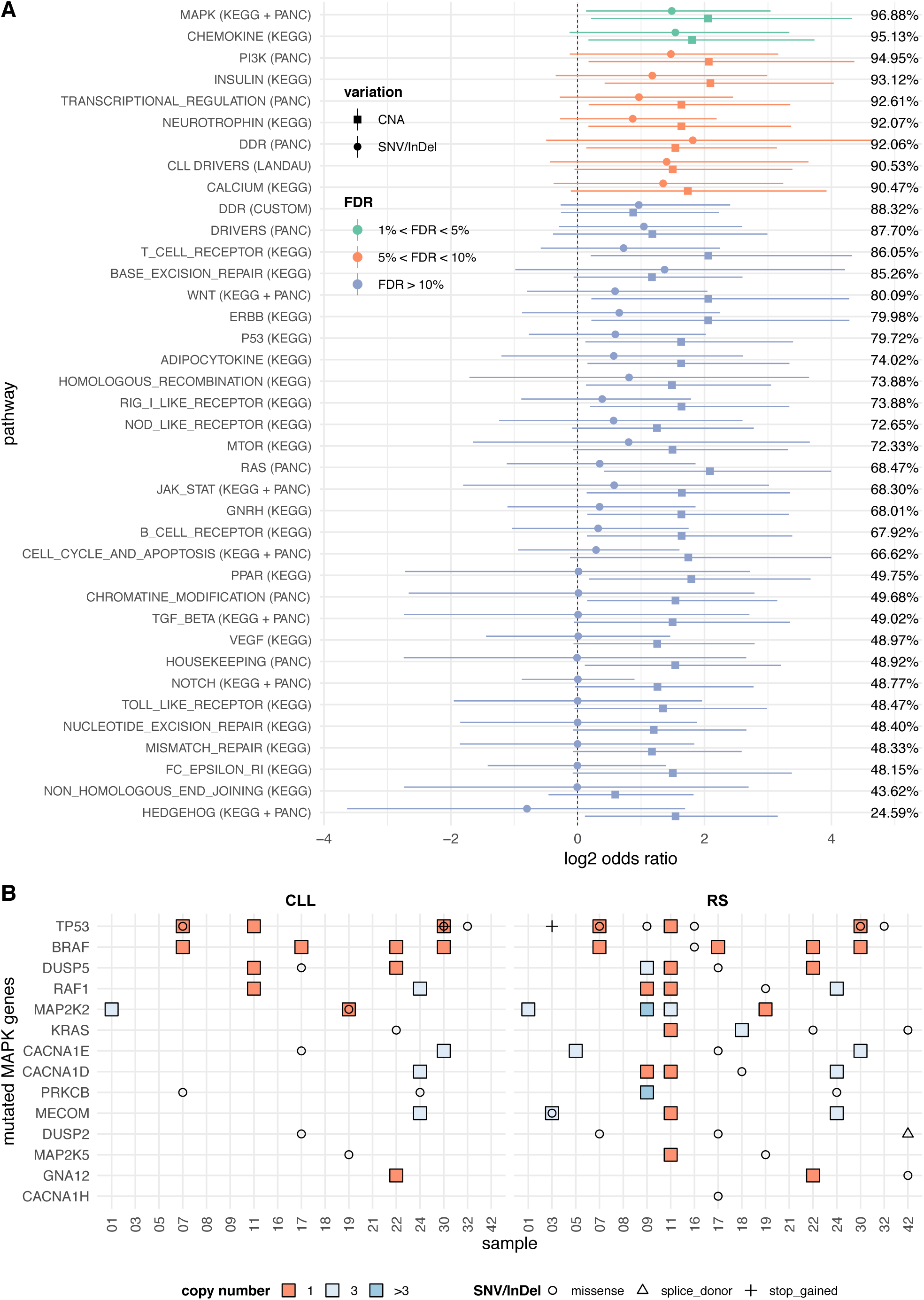
Pathway-based mutational burden analysis. A) For each pathway, we give the mean and 95% credible intervals of the ratio 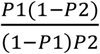 (in log_2_ scale), where P1 and P2 are the posterior probabilities that the RS and CLL phases, respectively, harbor an aberration. Separate estimates are given for SNVs/InDels (circles) and CNVs (squares). The percentages are the combined posterior probabilities that a particular pathway is more likely to harbor both a SNV/InDel and a CNV in the RS rather than the CLL phase. When this probability exceeds a threshold of 95%, which corresponds to FDR<5%, the pathway is considered significantly more mutated in the RS phase. B) Genes harboring SNVs, InDels and CNVs in the MAPK pathway. In total, 45 pathways were examined, but pathways appearing in both KEGG and PANC were merged resulting in 38 distinct pathways.

### Clonal evolution from CLL to RS

To further test the clinical and biological relevance of our findings, we wanted to understand whether any of the pathways mentioned above played a role in the clonal transition from CLL to RS. Therefore, we investigated clonal evolution patterns between CLL and RS in paired patient samples. Consistent with what is expected for a cohort of consecutively recruited patients, analysis of the IgHV locus demonstrated that all successfully tested RS cases carried related IgHV rearrangements implying that they originated from the same lymphoma stem cell (Supplementary Table 1). Only one case (08) was associated with EBV infection and was therefore likely to carry unrelated IgHV rearrangements due to an independent transformation event secondary to EBV.

For all 884 filtered somatic variants in our cohort, we estimated cancer cell fraction (CCF) values for all variants residing on autosomes (n=852, 96.4%; Supplementary Tables 2 and 3). Overall, we detected 186 subclonal and 145 clonal variants in the CLL phase and 294 subclonal and 211 clonal variants in the RS phase. A common pattern of clonal evolution observed in 10 out of 17 cases (58.8%) was characterized by clones in RS, which were present as either subclones (n=45 variants; e.g. cases 32 and 42 in Supplementary Figure 5) or clones in CLL (n=82 variants; e.g. case 01 in Supplementary Figure 5). More generally, we observed in all cases a degree of clonal expansion or contraction (i.e. a CCF increase or decrease in the transition from CLL to RS), or clonal stability (i.e. CCF values remain roughly the same between the two phases). In three cases (cases 03, 08 and 09), these transitions involved mutations that were mutually exclusive in CLL and RS samples (which explains the absence of connecting lines between them in Supplementary Figure 5). In case 08, very few mutations were detected in either CLL or RS consistent with the alternative transformation mechanism of EBV infection for this patient (Figure 2A, Supplementary Figure 5 and Supplementary Table 3).

The 211 clonal variants detected in RS were to a large extent not detectable at all in the corresponding CLL sample (n=84; 39.8%), hence they are putative contributors to the malignant transformation from CLL to RS. The *SVIL* and *DUSP2* genes showed this pattern in two recurrent samples each with clonal variants evolving at the time of transformation. In addition, *TP53* demonstrated both patterns of clonal evolution, each in two samples, with mutations that were either absent or subclonal in CLL evolving into clonal in RS (Supplementary Table 3).

Next, we performed an analysis of clonal evolution across different pathways, which demonstrated a high expansion probability for clones containing mutations in MAPK and transcriptional regulation genes, as well as mutations in DDR genes (Figure 4A). Among the MAPK pathway genes, purity-corrected CCFs of *TP53, DUSP2, KRAS, BRAF, CACNA1D, CACNA1H, GNA12, MECOM* and *RAF1* genes significantly increased during transformation from CLL to RS (Figure 4B). Some subjects harbored more than one MAPK pathway mutation in CLL and in some cases (03, 16 and 42) more than one expanding mutation was detected, while in other cases (17, 19 and 30) one particular mutation expanded, whereas the others contracted or remained relatively stable during the malignant transformation (Figure 4B).

**Figure 4:**
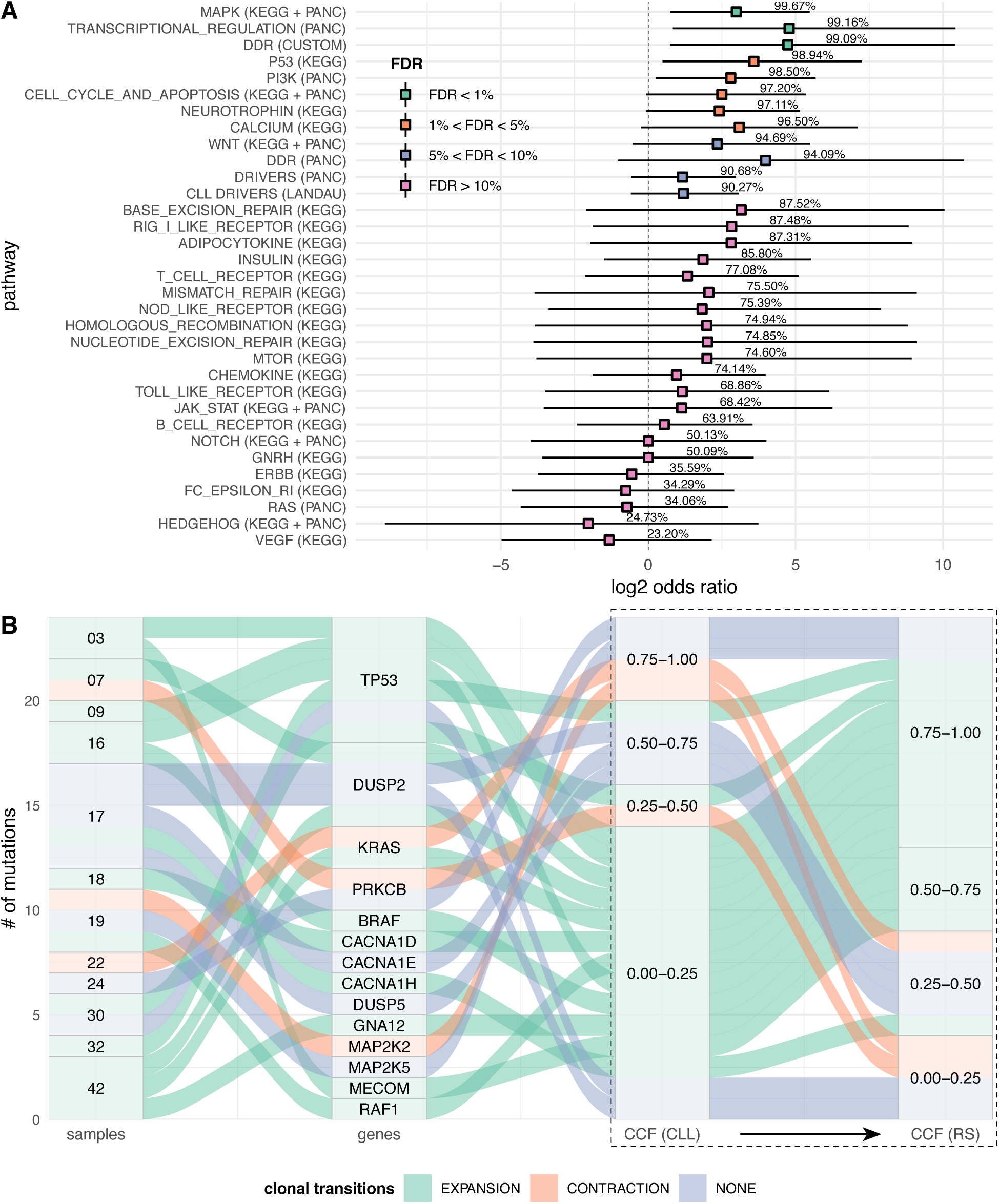
Pathway-based clonal analysis. A) For each pathway, we give the mean and 95% credible intervals of the ratio 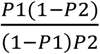 (in log_2_ scale), where P1 is the posterior probability that the pathway harbors a mutation that clonally expands (i.e. its CCF shows a significant increase) in the transition from CLL to RS. Similarly, P2 is the posterior probability that the pathway harbors a mutation that clonally contracts in the transition to RS. The percentages give the posterior probability that an expansion is more likely than a contraction, i.e. the probability that P1>P2. When this probability exceeds a threshold of 95%, which corresponds to FDR<5%, the pathway is considered highly likely to show clonal expansion rather than contraction. B) Alluvial plot summarizing clonal transitions (expansion, contraction or none) in the 14 genes of the MAPK pathway harboring SNVs or InDels. The part of the plot inside the dotted box directly illustrates the number of mutations expanding or contracting in the transformation from CLL to RS and their distribution across four different bands of CCF values. The remaining of the plot illustrates the distribution of expanding/contracting mutations across samples and genes.

### Non-coding variants

Furthermore, we explored the landscape of non-coding mutations during transformation. Considering only non-coding SNVs that were predicted to be functionally active (see Methods), we found a significantly higher number of mutations in the RS phase compared to the CLL phase (P=0.007; one-sided Wilcoxon signed rank test with continuity correction; Supplementary Figure 6). One sample had the known NOTCH1 3’UTR mutation in both CLL and RS (Supplementary Table 9). Interestingly, five patients carried mutation clusters in a region previously characterized as a functionally active PAX5 enhancer in CLL^19^. Three of these occurred in the RS phase only (Figure 5). When expanding this analysis to all filtered mutations in the entire PAX5 enhancer region, we found ten samples, each with 1-8 mutations, of which six had mutations only in the RS phase (Supplementary Table 9).

**Figure 5:**
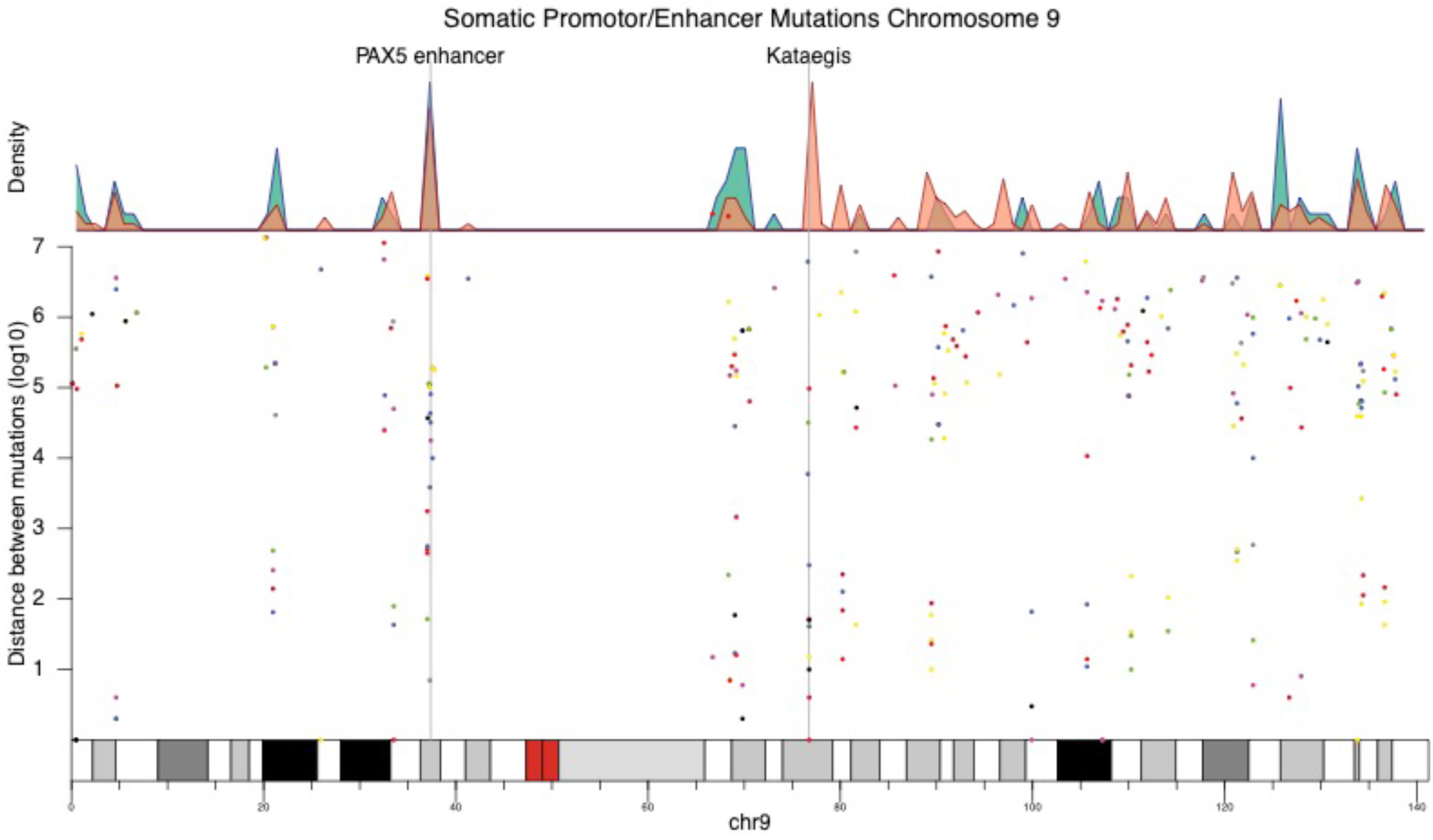
Kataegis regions in functionally active non-coding sites (see Supplementary Methods) across chromosome 9. The density plot shows the mutation spike at the PAX5 enhancer seen in both the CLL and RS phases, plus a kataegis region found only in the RS phase and located in close proximity to the RORB gene.

Using the functionally active non-coding SNVs to define regions of hypermutation, we found one kataegis region shared between the CLL and RS phase, 4 kataegis regions unique to the CLL phase and 103 unique to the RS phase (Supplementary Table 10). We examined the top three kataegis regions with the highest mutation burden in both CLL and RS (Supplementary Table 11). This showed that chromosome 9 contains an active non-coding kataegis region that is only present in RS and that is situated in close proximity to the gene RORB, which has been linked to multiple cancer types (Figure 5). We also found a kataegis region on chromosome 11 in the RS samples, which is topographically linked to CD44, and it is expected to contain an enhancer (Supplementary Figure 7A).

Mutation signatures were calculated from both exonic data (Supplementary Figure 7B) and functional non-coding data (Supplementary Figure 7C). Signatures were found in the functional non-coding data that were present in CLL or RS samples only. However, all three of these were signatures with an unknown function (Signatures 8, 12 and 16).

### Investigation of the transcriptome – confirmation of pathways

As a prelude to future functional analyses, we examined whether genomic aberrations seen at the DNA level (Figure 3) were mirrored by changes in gene expression. To address this question, we performed RNA expression profiling on 31 RS and 13 CLL lymph node samples using the NanoString PanCancer Pathways Panel (PANC), which includes 770 genes representing 13 canonical cancer pathways involved in various cellular processes, such as cell cycle regulation, apoptosis, proliferation and differentiation, plus 30 genes selected based on results of the WGS and previous RS reports (see Supplementary Table 5).

We found 22 down-regulated and 21 up-regulated genes in RS compared to CLL (Figure 6A). Twenty-three of these have been reported in CLL or other lymphomas^26^. The remaining differentially expressed genes (n=20) were down-regulated in RS and they are known tumor-suppressor genes (e.g. *TSC1, STK11, IKBKB, PIK3R1*). The majority of up-regulated genes in RS are known oncogenes (e.g. *GRB2, HSP90, CDK4*) (Figure 6A). Pathway enrichment analysis showed that the PI3K, JAK-STAT and P53 pathways are the most likely to harbor differentially expressed genes (FDR<5%) (Figure 6B). Moreover, five pathways (PI3K, cell cycle and apoptosis, RAS, PANC Drivers and MAPK) harbored 10 or more differentially expressed genes (Figure 6C).

**Figure 6:**
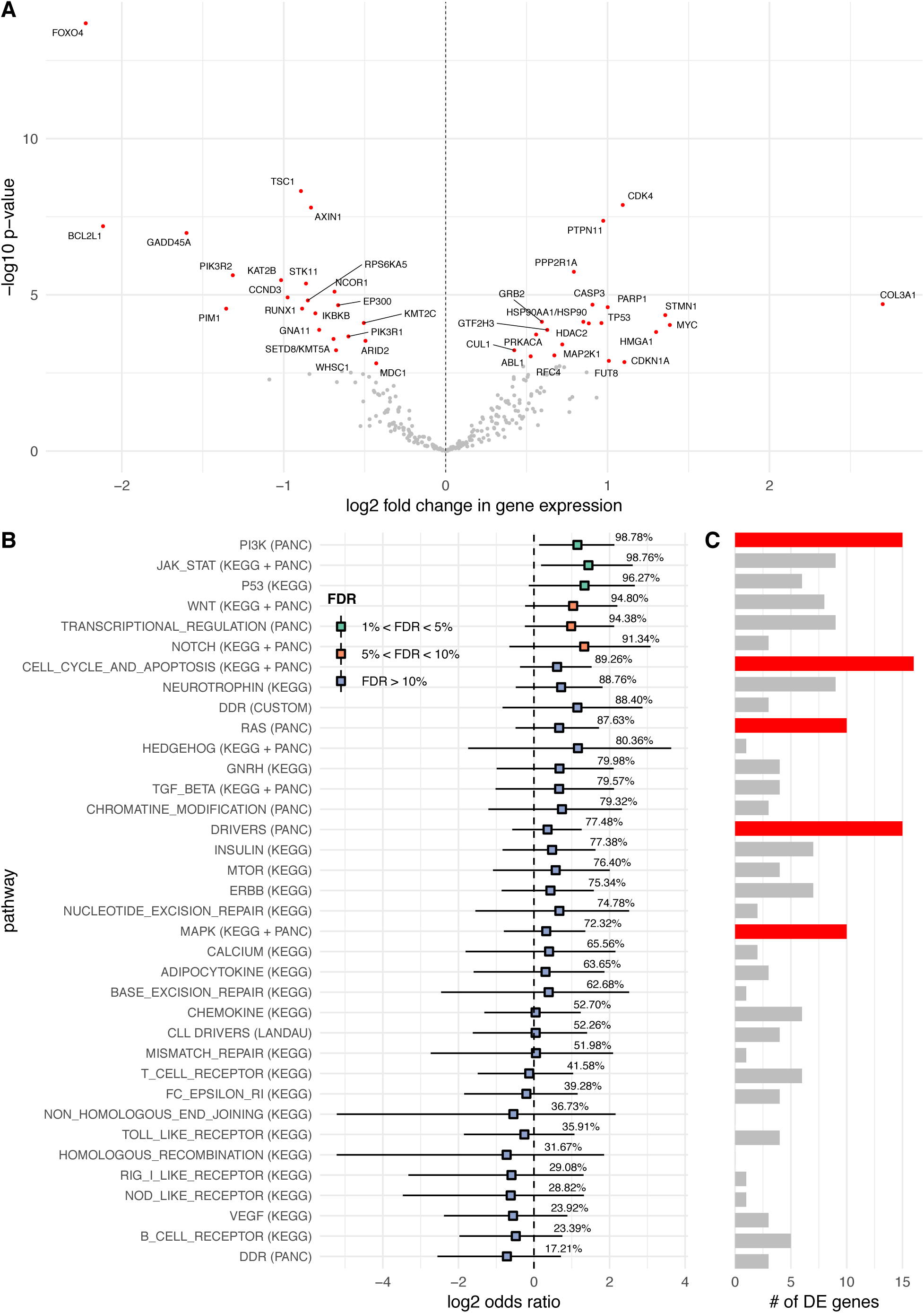
Differential expression and enrichment analysis. A) Volcano plot indicating up- and down-regulated genes in RS when compared to the CLL phase. B) For each pathway, we give the mean and 95% credible intervals of the ratio 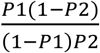 (in log_2_ scale), where P1 and P2 are the expected fractions of differentially expressed genes in the pathway and not in the pathway, respectively. Each percentage indicates the posterior probability that P1>P2. When this probability exceeds a threshold of 95%, which corresponds to an FDR<5%, we say that the pathway is enriched in differentially expressed genes. C) Absolute number of differentially expressed genes in each pathway.

## Discussion

During the last decade, significant advances in the treatment of CLL have translated into greatly improved clinical outcomes. However, CLL still remains largely incurable, and high-grade transformation, most commonly seen in heavily pre-treated patients with CLL, remains an area of high unmet clinical need with a dismal outcome. Due to the rarity and aggressive nature of RS, clinical trials of novel agents are difficult to perform, and there is a lack of suitable disease models for rational drug design. Contrary to other rare cancers characterized by a specific single genetic aberration, the molecular profile of RS is highly heterogeneous^3,15,16,19^ making genomic analysis challenging.

Here, we show results from a novel approach of integrative WGS analysis of paired CLL and RS samples from a cohort of patients recruited into a frontline clinical trial. We use a combination of coding and non-coding SNVs, InDels and CNAs, as well as RNA expression profiling of 800 cancer-related genes, to interrogate specific genes and pathways involved in the evolution of RS.

We confirm the frequent presence of acquired pathogenic SNVs in *TP53*, the recurrent indels in the PEST domain of *NOTCH1*, acquired CNAs in *CDK2NA*, and *MYC* deregulation (over-expression) in the RS phase identified through differential RNA expression analysis. Moreover, we identify activating mutations and amplification in XPO1 and copy number losses of *TRAF3* as additional genes frequently targeted by mutations in RS. *TRAF3* is a negative regulator of the non-canonical *NFκB* signaling pathway. Deletions of *TRAF3* lead to overexpression of *NFκB*-inducing kinase (NIK). Inhibitors of NIK belong to a promising new class of drugs entering clinical development^29^. Exportin 1 (XPO1) belongs to a family of proteins providing cytoplasmic-nuclear transport for a large number of cargo molecules. Activating mutations and copy number gains in *XPO1* have been reported in both hematologic and solid neoplasms^18,23,30,31^. The XPO1 inhibitor Selinexor has recently been evaluated in a phase 1 trial of patients with relapsed or refractory non-Hodgkin lymphomas demonstrating acceptable safety and response (CR and PR) in 35% of patients, including patients with RS^32^.

Four CLL driver genes were recurrently mutated in the RS phase only, suggesting a role as clonal drivers: *PTPN11*, a positive regulator of MAPK-RAS-ERK signaling pathway^33,34^; *MGA*, a repressor of MYC function known to be mutually exclusive to *MYC* genetic lesions in RS; the B-cell transcription factor *IKZF3* ^35,36^ and *BAZ2A* regulated by the microRNAs MIR15a/16-1^37^, which are commonly deleted in CLL. Importantly, our extended analysis of 55 patients shows that SNVs/InDels in *MGA* are one of the most common findings in RS.

Our genome-wide approach allowed us to associate mutations in targetable pathways that have not previously been implicated in RS transformation. We show that the MAPK pathway has a higher somatic SNVs/InDels and CNAs burden in RS compared to the CLL phase. Clones containing MAPK pathway mutations demonstrate high expansion probability. The MAPK pathway was also one of the pathways with the highest number of differentially expressed genes between RS and CLL phases.

Targeted treatments with RAF and MEK inhibitors are approved for clinical use in malignant melanoma^38^. In B-cells, MAPK signaling is initiated downstream to BCR activation and therapeutic targeting of BCR signaling through BTK, PI3K and SYK inhibition has a suppressive effect on MAPK signalling^39^.

In CLL, mutations in MAPK pathway genes have been reported at a frequency of 8% with an accumulation in poor-prognosis cases^23,40^ and treatment failure on BTK inhibitors is associated with the emergence of RS. Treatment of primary CLL cells *in vitro* with MEK inhibitors resulted in induction of apoptosis, suggesting potential activity against CLL^40,41^. The effect of MEK inhibition on RS has not been studied and there are no clinical trials studying MAPK pathway inhibition in CLL or RS.

Finally, we extend previous observations of the functional significance of non-coding mutations in cancer. A hypermutated region 330kb upstream of the *PAX5* gene was previously described as an enhancer region of *PAX5*. In this cohort, which was enriched for good prognosis CLL, mutations were seen more frequently in patients with hypermutated IgVH and del13q abnormalities. The authors therefore hypothesized that these *PAX5* enhancer mutations are implicated in driving early CLL development. Interestingly, the same enhancer region was also mutated in 29% of DLBCL controls. Here, we show that non-coding mutations in *PAX5* enhancer elements are also potential drivers of transformation, as they occur preferentially in RS.

In conclusion, we show that integrated WGS combined with RNA expression profiling identifies multiple potential therapeutic targets for clinical evaluation. A UK Phase 2 adaptive clinical platform trial, The STELLAR study (2017-004401-40), is now open to recruitment for patients with RS to evaluate the novel therapies identified in this study.

## Methods

### Samples acquisition

Peripheral blood (CLL), tumor (RS; Formalin-Fixed Paraffin-Embedded blocks, FFPE) and germline (GL; from saliva) triplet samples were available from 17 of the 37 evaluable patients enrolled in the CHOP-OR trial^21^. Ethics approval for CHOP-OR was obtained from the National Research Ethics Service Committee South Central – Oxford A (REC reference number: 10/H0604/85).

An independent cohort of patients diagnosed with RS between 2005 and 2016 by the Department of Pathology, Skåne University Hospital, Lund University, Sweden underwent DNA sequencing (n=21) and RNA expression profiling (n=21) (ethics approval: Southern Sweden, reference number: 2016/1054). Moreover, DNA from RS tumors was available from 7 additional patients from the CHOP-OR study and from 10 patients with biopsy-confirmed RS treated outside of the clinical trial (ethics approval: Hematology Collection Protocol, HTA License Number 12217, Oxfordshire C, REC: 09/H0606/5). See Figure 1.

### Purification of DNA and RNA

#### CHOP-OR cohort

Paraffin was removed from FFPE 10μm sections scraped from 5-10 slides using the Adaptive Focused Acoustics (AFA™) on the M220 Focused-ultrasonicator. DNA was thereafter purified with the truXTRAC FFPE DNA Kit according to the manufacturer’s protocol (Covaris Inc., Woburn, MA, USA), but using optimized reverse cross-linking conditions as previously described^22^. Peripheral blood samples were subjected to a ficoll gradient centrifugation to isolate mononuclear cells. Percentage of CD19+ cell was verified by flow cytometry and samples with a purity <70% were purified using magnetic beads according to the manufacturer’s protocol (Miltenyi Biotec, Bergisch Gladbach, Germany). Extraction of genomic DNA was done using the QIAamp Mini kit according to the manufacturer’s protocol (Qiagen, Hilden, Gemany). For the germlines, DNA was extracted from saliva using the prepIT•L2P according to the manufacturer’s protocol (DNA Genotek, Ottawa, Ontario, Canada). RNA was extracted from 11 FFPE blocks using the protocol specified below.

#### Independent cohort

FFPE blocks were prepared in 10μm sections, transferred to 1.5 mL tubes (60-100 μm per tube) and in order to prevent RNA degradation stored in −80 degrees C immediately after the preparation. 46 μm of FFPE tissue was used for combined DNA and RNA purification. After removing the paraffin from the FFPE sections using Qiagen’s Deparaffinization Solution (Qiagen, Hilden, Germany), total RNA and genomic DNA were purified using the AllPrep® DNA/RNA FFPE Kit according to the manufacturer’s protocol (Qiagen, Hilden, Germany). DNA was obtained from 21 samples and RNA extraction was successful in 21 of 23 Lund University cohort samples.

DNA from all cohorts was quantified using the Qubit® 2.0 Fluorometer (Thermo Fisher Scientific, Waltham, MA, USA). DNA and RNA quantity and fragment length were analyzed using the Agilent Bioanalyzer (Agilent, Santa Clara, CA, USA).

### Whole Genome Sequencing

WGS libraries were prepared using optimized Illumina protocols depending on DNA type (Illumina, San Diego, California, United States). The Early Access FFPE-extracted gDNA Library preparation kit (Illumina) was used for FFPE-derived DNA. In this protocol, FFPE DNA was first treated with the FFPE DNA Restoration kit (Illumina) in order to repair damaged DNA and to achieve high-quality DNA for further WGS library preparation. The TruSeq DNA HT Sample Prep Kit (PCR-free) or TruSeq Nano DNA LT Library Prep Kit (with PCR amplification) were used, both according to the manufacturer’s protocol, for CLL and germline DNA library preparation depending on the amount of DNA available (Illumina, San Diego, California, United States).

Libraries were subjected to 2×100 bp paired-end sequencing on a HiSeq 2500 or 2×150 bp a HiSeq 4000 instrument (Illumina, San Diego, California, United States), to a mean sequencing depth of 87X for CLL (range 34X-129X), 88X for RS (range 42X-132X), and 44X for germline (range 25X–67X) samples.

### Bioinformatics analysis of WGS data

Raw reads from each triplet of germline (GL), blood (CLL) and lymphoma (RS) samples for each patient were aligned against the hg19 human reference genome (release GRCh37) using Illumina’s Whole Genome Sequencing workflow v4.0.0, and somatic SNVs and InDels were called for each GL/CLL and GL/RS pair using Illumina’s Tumor-Normal workflow v1.1.0. Somatic variants with PASS filter, read depth DP≥10, and VAF≥5% in the CLL phase or VAF≥10% in the RS phase were retained, and they were annotated using Ensembl’s Variant Annotation Predictor v90, which included functional predictions from SIFT and PolyPhen. Variants which were predicted as having HIGH impact by VEP, or which were flagged as deleterious by SIFT or probably damaging by PolyPhen were kept for further analysis. Copy number analyses were carried out as described in Schuh et al (2018)^42^.

### Targeted Sequencing of independent cohorts

A targeted xGen® Lockdown® Probes panel (Integrated DNA Technologies, Inc., Skokie, IL, USA) and a TruSeq Custom Amplicon (TSCA, Illumina, Inc., San Diego, CA, USA) panel were designed to target 56 and 28 genes and hotspots, respectively, of specific relevance to aggressive CLL, Richter’s transformation, and other genes of interest. Gene selection was based on findings in the WGS of the training cohort (CHOP-OR) and/or on the available literature. Targets are specified in Supplementary Table 12. Sequencing was performed using the MiSeq platform (Illumina, Inc., San Diego, CA, USA) 2×150bp paired-end sequencing.

### Bioinformatics analysis of TGS data

For the 56-genes panel, alignment was performed using BWA-mem v0.7.10 against the hg19 human reference genome (reference GRCh37) and deduplication and error correction were performed by Connor v0.5.1 (https://github.com/umich-brcf-bioinf/Connor) changing the default min_family_size_threshold to 1. Variant calling was performed using Platypus v0.8.1^43^.

For the 28-genes panel, the TSCA workflow MiSeq Somatic Variant Caller v3.2.3 (Illumina) was used to perform initial alignment against the same human reference genome and variant calling using the default settings. Following this, a second alignment using BWA-mem v0.7.12-r1039 and indel realignment with GATK v3.5 were used before variant calling using Platypus v0.8.1^43^ in order to detect additional variants.

For both panels, functional sequence variants were identified by filtering on quality, functional consequence as determined by VEP v84, and predicted pathogenicity by *in silico* tools (SIFT and PolyPhen), previous published literature, or presence in the COSMIC database v73. Variants reported in ≥5% of general population or reported as germline in ClinVar were removed. All variants were confirmed by visual inspection in IGV^44,45^.

### IgHV mutational status

IgHV mutational status was determined by Sanger and targeted next-generation sequencing as previously described^46^. IgHV analysis was performed on CLL and RS tumor samples in order to determine their clonal relatedness. However, the analysis was conclusive in only eight cases. Therefore, comparative analysis of clonally related versus unrelated RS cases was not possible to perform (see Supplementary Table1).

### Annotation and processing of non-coding regions

Non-coding annotation was performed by intersection with sites of expected non-coding functionality in CLL. Functional non-coding sites were determined as follows: the region must be within a peak of ATAC-seq activity in 20% of CLL samples (>21/106) from Beekman et al. (2018)^47^. Additionally, the region must be identified as either active promotor, strong enhancer1, or strong enhancer2 by CHROMHMM^48^ in 3 or more samples including 7 CLL and a further 15 from various blood cells (2x csMBC, 1x ncsMBC, 3x GCBC, 3x NBCB, 3x NBCT and 3x PCT). Briefly, CHROMHMM was used to combine 6 histone modification marks in each CLL sample to identify genomic functionally active regions, as described in Beekman et al. (2018)^47^. Association with TAD (Topographically Associated Domains) was carried out using CLL and B-cell Hi-C, ATAC-seq and H3k27ac histone modification marks. Using the combination of histone modification marks and ATAC-seq together is expected to increase the proportion of mutations of functional impact caused by AID. Kataegis was identified according to Lawrence et al. (2013)^49^ using the pooled functional non-coding data, and plots were produced with the R library KaryoploteR^50^. Mutation signatures were analyzed on the same data based on the methods of Alexandrov et al. (2013)^51^, using the R package deconstructSigs^52^.

### Digital-multiplexed Gene Expression Profiling

To assess gene expression differences in tumor tissue from CLL lymph nodes versus transformed Richter’s DLBCL we applied the NanoString PanCancer Pathways Panel (NanoString Technologies, Seattle, WA, USA) including 770 genes representing 14 canonical cancer pathways and 30 custom-selected genes (in total 800 genes) selected based on findings in the initial WGS and also based on potential importance to RS, as presented in previous publications (Supplementary Table 5). In total 13 CLL lymph node samples (Lund University cohort) and 31 Richter’s DLBCL samples were analyzed (11 CHOP-OR RS samples and 20 Lund University RS samples). After purification, RNA was quantified and quality-assessed with the Agilent Bioanalyzer (Agilent, Santa Clara, CA, USA). All samples were controlled for degree of RNA integrity and also for fragment length using the smear analysis function in the Agilent Bioanalyzer software. To correct for suboptimal RNA integrity, based on the percentage of degraded RNA with a fragment length falling between 50-300nt, a target input of 140 ng was calculated using the following formula:

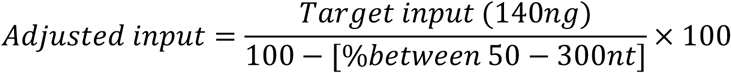

RNA was hybridized for 16 hours at 65° C with the PanCancer Pathway Code Set and the spike-in custom CodeSet including the additional 30 genes of interest (NanoString Technologies, Seattle, WA, USA) (Supplementary Table 2). Purification, binding and scanning of the hybridized probes to the cartridge was performed on the nCounter® SPRINT Profiler. Raw data files from the nCounter® SPRINT Profiler were imported into the nSolver 4.0.62 Analysis software (NanoString Technologies, Seattle, WA, USA) and were checked for data quality using default quality control settings.

### Statistical analysis

All statistical analyses were performed and graphics were generated using R v3.5.1^53^. A detailed mathematical description of the various statistical analyses we used is given in the Supplementary Material. Below, we give a general overview of these analyses.

First, we examined whether the CLL and RS phases were differentially mutated. In order to increase the power of our analysis, instead of focusing on individual variants or on individual genes, we focused on sets of genes (i.e. pathways). These included all KEGG signalling, cell cycle, apoptosis, and DNA damage repair (DDR) pathways, all NanoString PanCancer pathways, a set of 44 CLL driver genes, and a custom set of 21 DDR genes previously described in solid tumours. In total, we examined 45 different gene sets/pathways. By adopting a Bayesian approach, we calculated, for each pathway and each phase (CLL or RS), the posterior probability (i.e. the probability given the data) that the phase harbours both a somatic SNV/InDel and a CN event. The pathways that were more likely to harbour genomic aberrations in the RS rather than in the CLL phase with probability higher than 95% (FDR<5%) were identified as differentially mutated, and they were retained for further analysis.

In a second stage, we examined the clonal structure of each tumour sample, using a previously published statistical approach^54,55^. Briefly, we used a Bayesian non-parametric clustering methodology, which considers the purity of each sample, as well as the multiplicity and VAF of each somatic variant, and it estimates a cancer cell fraction (CCF) for each. Variants with CCF>0.85 with probability higher than 95% were considered clonal, those with CCF<0.85 with equally high probability were consider sub-clonal, while the rest were deemed of uncertain clonality. Somatic variants demonstrating an increase in CCF from CLL to RS larger than 25% were deemed indicative of a clonal expansion event, while those with a CCF decrease of equal magnitude were considered evidence of clonal contraction. By examining the clonal expansion and contraction events in each pathway, we identified those pathways that had more than 95% probability (FDR<5%) of harbouring clonal expansion instead of clonal contraction events in the transition from CLL to the RS phase.

Finally, we performed differential gene expression analysis between 13 CLL and 31 RS samples. Only genes with at least 100 counts per million in each sample were analysed, resulting in a dataset with 301 genes. edgeR v3.24.0^56^ was used for normalisation and for testing for differential expression using likelihood ratio tests. Genes with FDR<1% were identified as differentially expressed, and they were further tested for enrichment against the 45 pathways mentioned above.

## Supporting information

HICF2 Consortium members

## Acknowledgments

We acknowledge the contribution to this study made by the Oxford Centre for Histopathology Research and the Oxford Radcliffe Biobank, which are supported by the NIHR Oxford Biomedical Research Centre. JK was supported by research grants from the Tegger Foundation, the Gunnar Nilsson Cancer Foundation, Blodsjukas förening i Södra sjukvårdsregionen, Stiftelsen Siv-Inger och Per-Erik Anderssons minnesfond, the Royal Swedish Academy and the Swedish Medical Association. This study was partly funded by the National Institute for Health Research Oxford Biomedical Research Centre. This publication presents independent research commissioned by the Health Innovation Challenge Fund (R6-388 / WT 100127), a parallel funding partnership between the Wellcome Trust and the Department of Health. This research was also supported by by the Wellcome Trust Core Award (203141/Z/16/Z). The views expressed in this publication are those of the authors and not necessarily those of the National Institute for Health Research, the UK National Health Service, the UK Department of Health, the University of Oxford or the Wellcome Trust.

## Supplementary Tables

**Supplementary Table 1:** Clinical characteristics of the CHOP-OR cohort (n=17) at the time of inclusion (n=no of assessed subjects; N=subjects within the group with complete data). The second spreadsheet shows IGHV status of the CHOP-OR cohort.

**Supplementary Table 2:** Filtered SNVs and InDels in each sample and phase identified using whole genome sequencing.

**Supplementary Table 3:** Wide format of the data in Supplementary Table 4 showing the clonal transitions of each SNV/InDels from CLL to RS. Only mutations in autosomal chromosomes are shown, and only mutations for which the estimated CLONALITY is not UNCERTAIN.

**Supplementary Table 4:** CNVs in each sample and phase, as well as genes affected by each copy number event.

**Supplementary Table 5:** KEGG, PanCancer and custom gene sets used for the mutational burden, clonal structure and pathway enrichment analysis. We also show 30 genes of special interest supplementing the 770 genes of the NanoString PanCancer Pathways Panel used for digital multiplexed gene expression profiling.

**Supplementary Table 6:** Recurrent genes (n=33) in the 17 CHOP-OR samples as identified using whole genome sequencing. The 17 genes which are recurrent with respect to mutations that expand clonally, or which are found exclusively in the RS phase are indicated in red. The second spreadsheet shows the mutations (n=80) in these 33 recurrent genes. The seven mutations that show clonal expansion in the RS phase and which are also recurrent are indicated in red.

**Supplementary Table 7:** Mutation frequency of genes inferred using targeted sequencing. For the same genes, the mutation frequency from whole genome sequencing is also reported.

**Supplementary Table 8:** Filtered SNVs and InDels in each sample of the independent cohorts identified using targeted sequencing.

**Supplementary Table 9:** Distribution of non-coding mutations between CLL and RS phases in the NOTCH1 3’UTR and in a region previously characterized as a functionally active PAX5 enhancer in CLL.

**Supplementary Table 10:** Sites of somatic hypermutation (kataegis) with 6 or more mutations less that 2 standard deviations of the inter-mutational distance apart.

**Supplementary Table 11:** Top 3 kataegis regions with the greatest number of mutations (reported for both the CLL and RS phases). TAD – Topographically Associated Domain.

**Supplementary Table 12:** Targets for targeted sequencing on the Illumina MiSeq platform using an xGen® Lockdown® Probes panel and a TruSeq Custom Amplicon panel, covering 56 and 28 genes and hotspots, respectively.

## Supplementary Figures

**Supplementary Figure 1:**
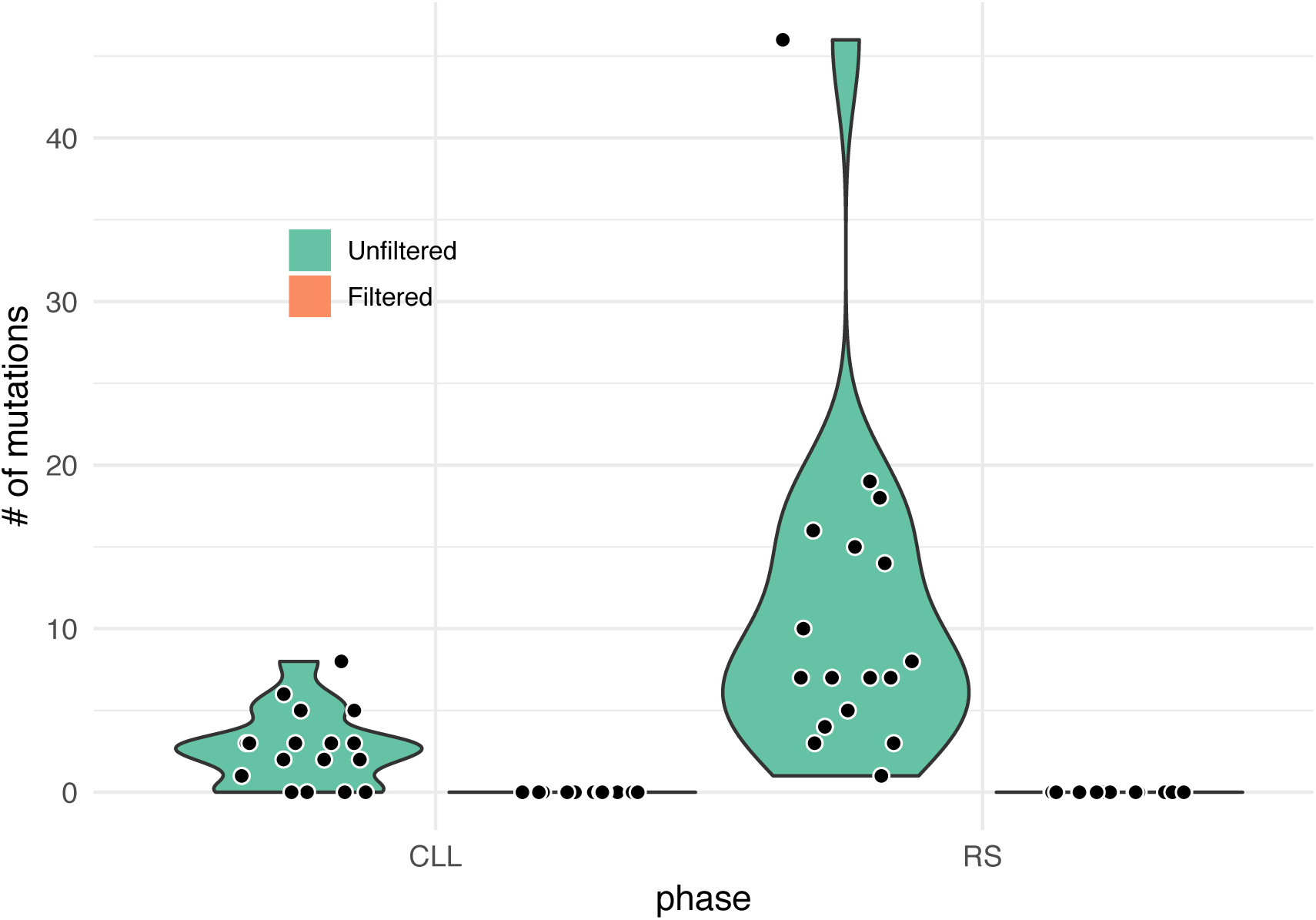
Number of mutations in the T-cell receptor-{C,V,J} gene locus in the CLL and RS phases across all samples before and after filtering.

**Supplementary Figure 2:**
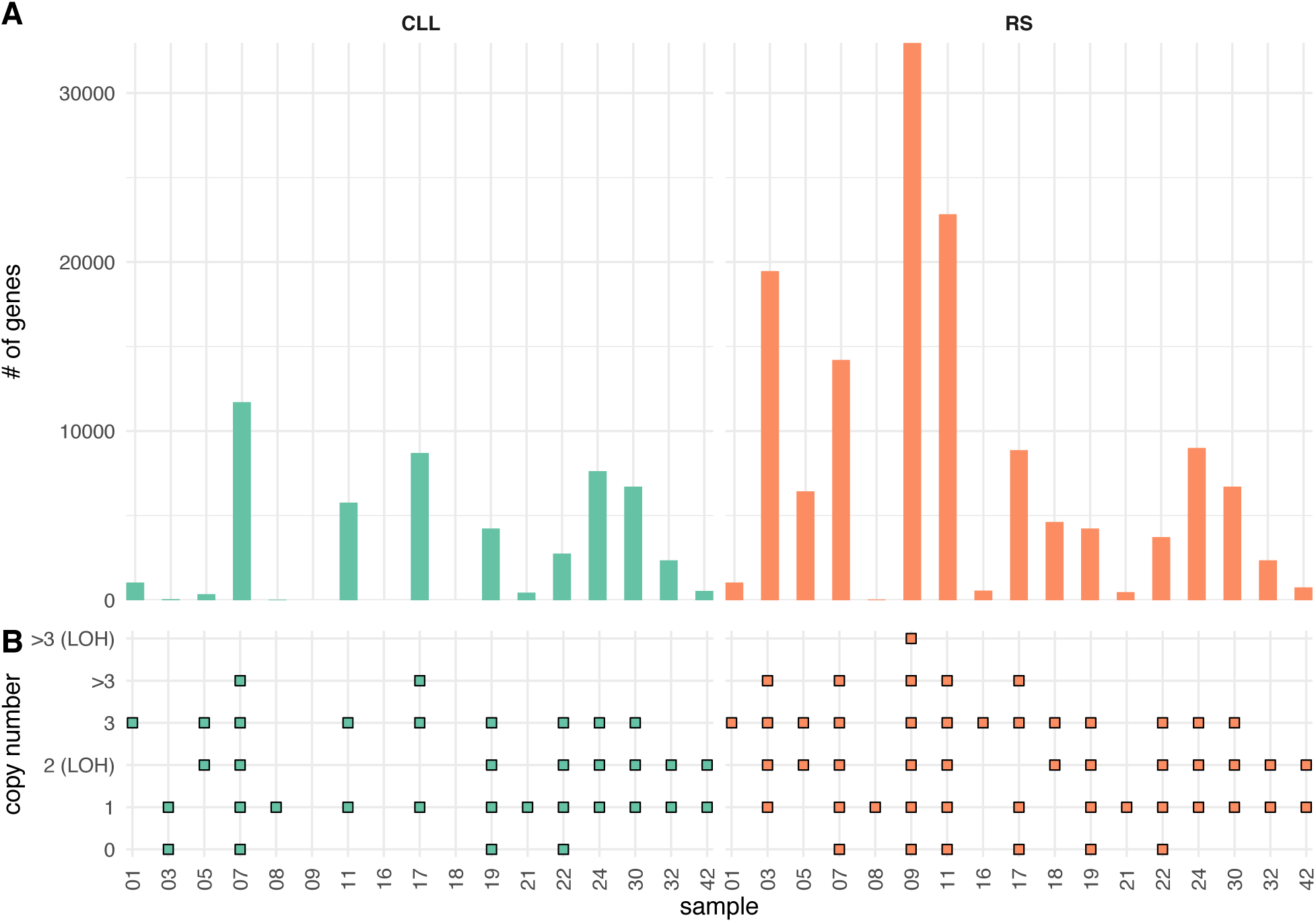
Overview of identified CNAs. A) Number of genes affected by CNAs in CLL and RS in each sample. B) Types of copy number events in each sample and phase.

**Supplementary Figure 3:**
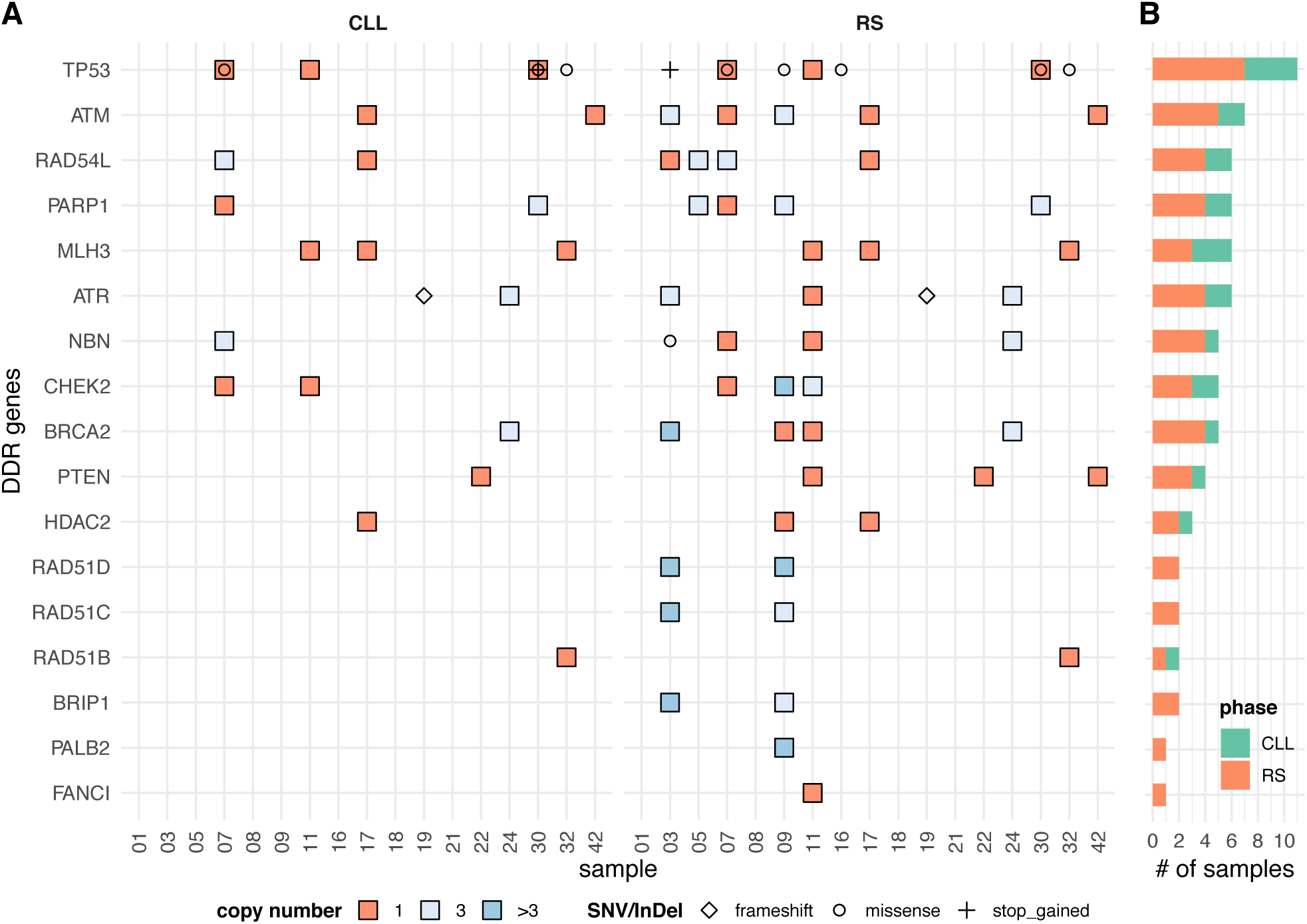
Genomic aberrations detected by whole genome sequencing in DDR genes. A) Distribution of SNVs, InDels and CNAs across samples and phases. B) Number of samples harboring a genomic aberration in each phase in each DDR gene. 21 genes were examined, but only 17 were found mutated.

**Supplementary Figure 4:**
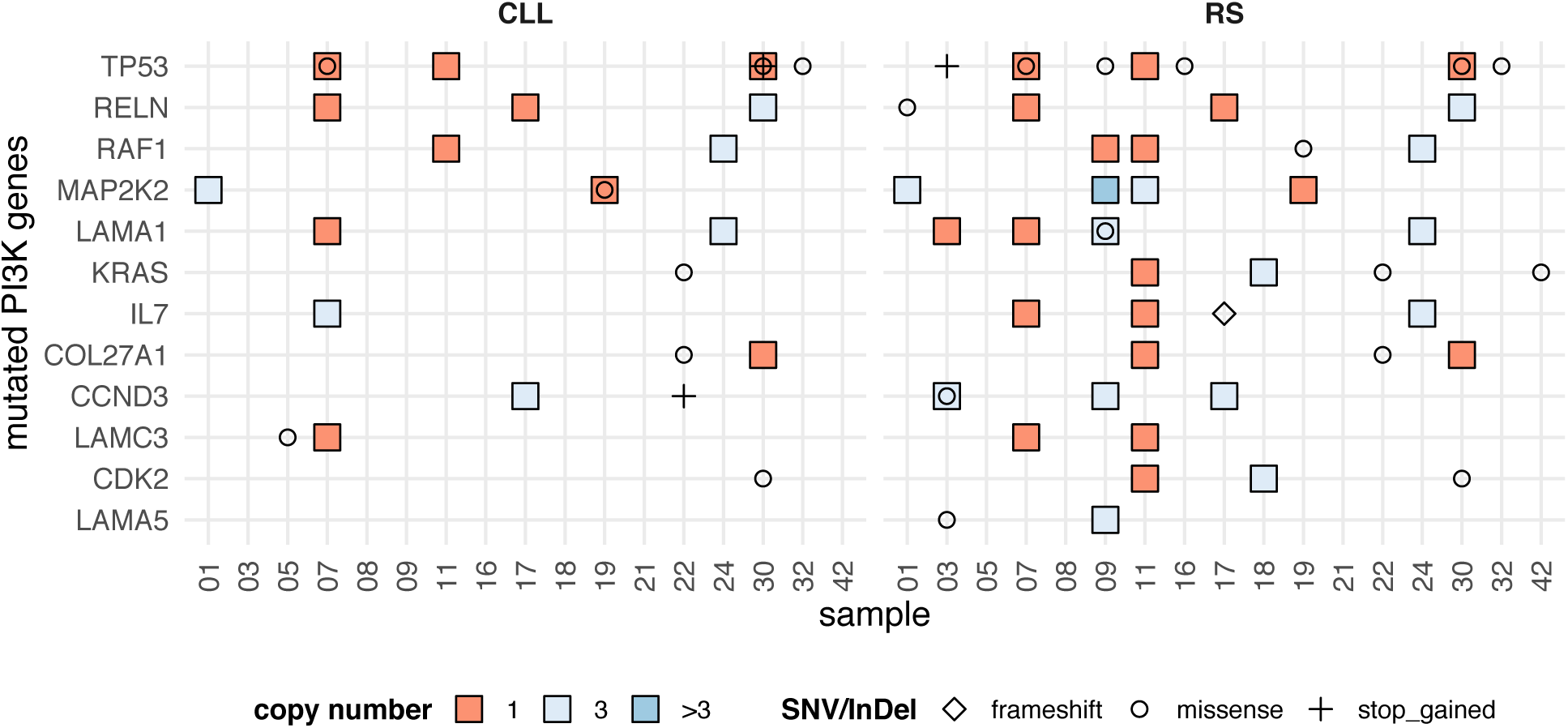
Genomic aberrations in the PI3K pathway. Only mutated genes (n=12) in the pathway are shown.

**Supplementary Figure 5:**
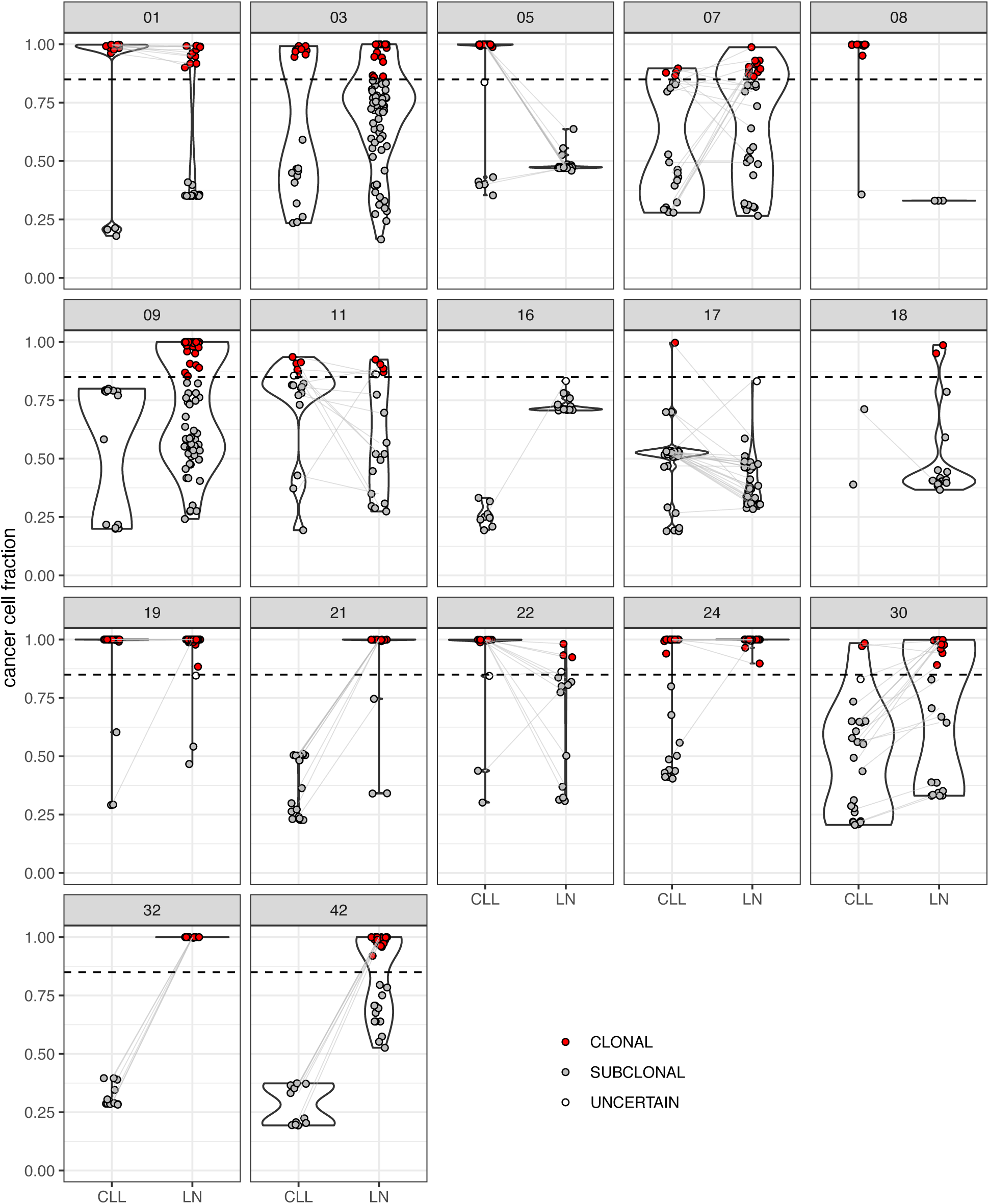
Transitions in the values of cancer cell fractions (CCF) of somatic mutations during transformation from CLL to RS in the 17 CHOP-OR cases. We observe clonal expansions (i.e. CCF values increase from CLL to RS), clonal contractions (i.e. CCF values decrease from CLL to RS) or clonal stability (i.e. CCF values remain roughly the same between CLL and RS). In extreme cases (e.g. cases 03, 08 and 09), CCF values increase from 0 in CLL or decrease to 0 in RS, which explains the absence of connecting lines between the CLL and RS phases in these cases.

**Supplementary Figure 6:**
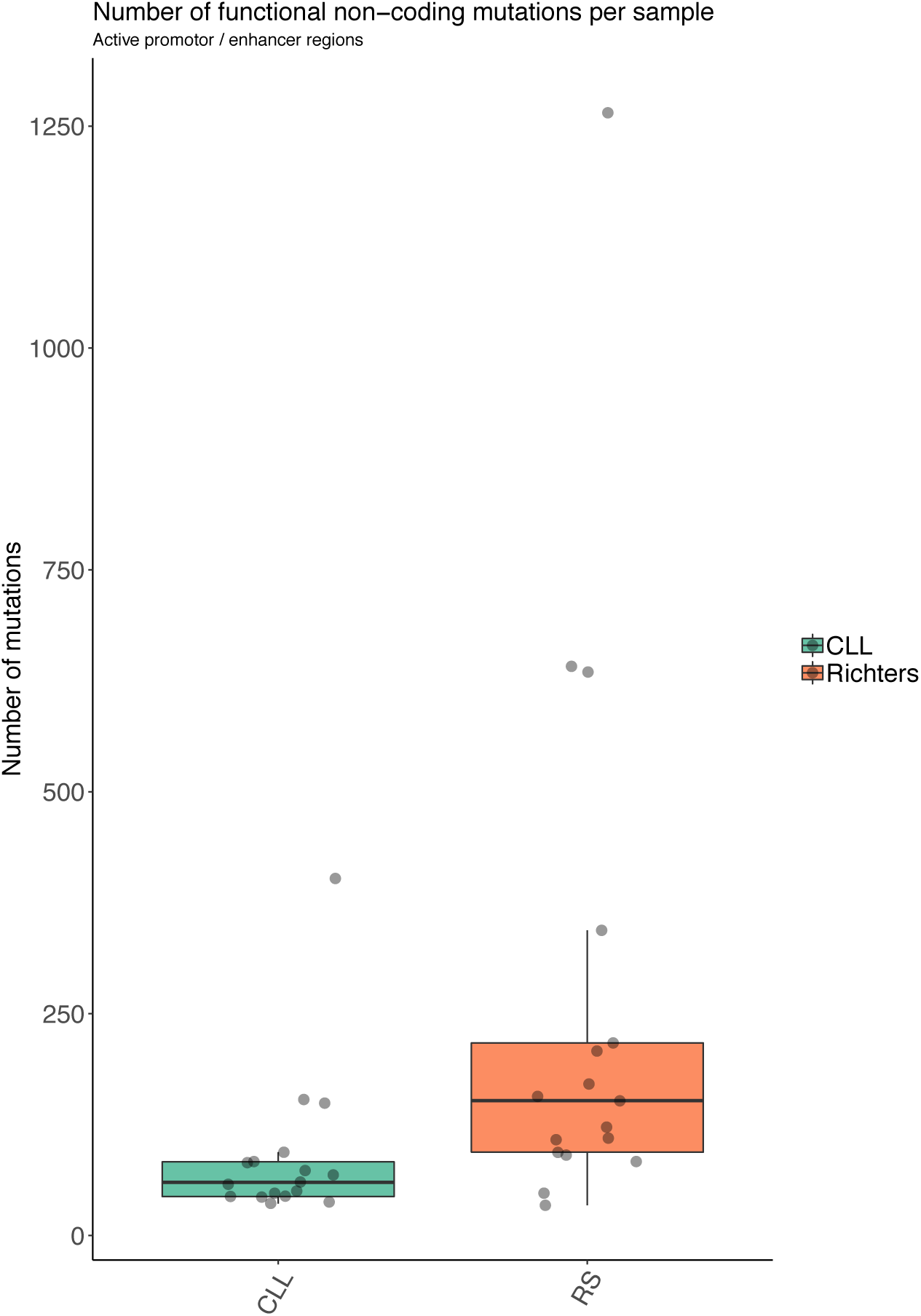
Numbers of functionally active non-coding SNVs in the CLL and RS phase across all samples, in predicted active promotor/strong enhancer regions. Average mutation number is found to be significantly more mutated in the RS phase (P=0.007; one-sided Wilcoxon signed rank test with continuity correction). The combination of histone modification marks and ATAC-seq is expected to minimize the effects of AID-mediated SNVs (Beekman et al. 2018).

**Supplementary Figure 7:**
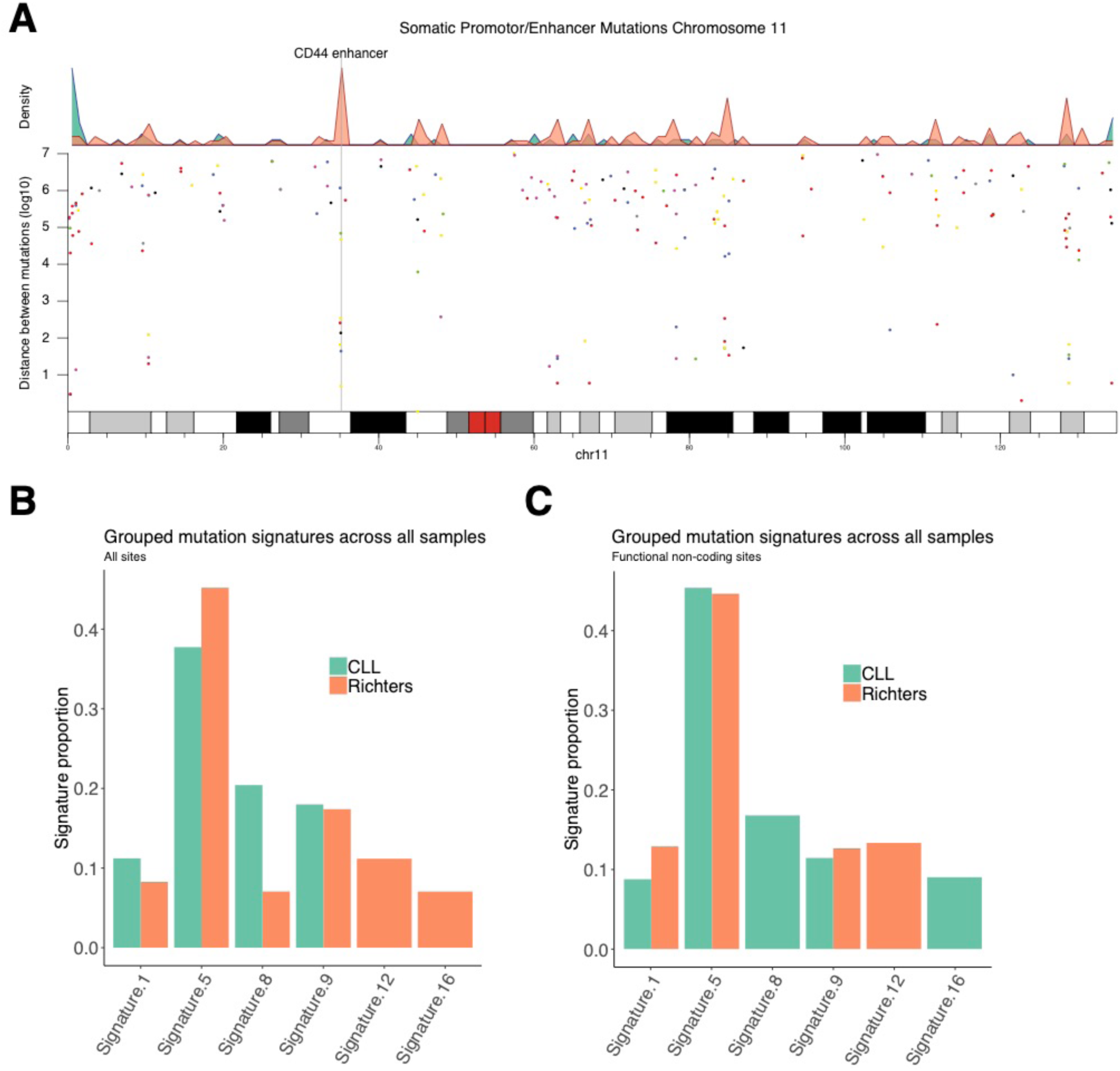
A) A kataegis region on chromosome 11 (Chr11:35,163,868-35,163,917) found in the RS samples. The region is topographically linked to CD44, and it is expected to contain an enhancer. B) We also show the proportion of mutation signatures (Alexandrov et al., 2013) in all exonic SNVs (n=884) across all samples, and C) in all functionally active non-coding SNVs (n=6004) across all samples. Signatures are as follows: Signature 1: ageing, Signature 5: ageing, Signature 8: unknown, Signature 9: AID, Signature 12: unknown, Signature 16: unknown.

## Supplementary Statistical Methods

In this supplementary material, we provide details of the statistical analyses applied in the main text. In all cases, posterior probabilities were estimated using self-adjusting Hamiltonian Monte Carlo[1], as implemented in `Stan`[2]. The exception to this was the estimation of cancer cell fraction (CCF) values in the section Clonal Analysis, where automatic differentiation variational inference (ADVI) was adopted for computational efficiency[3] (again using `Stan`).

### 1. Mutational burden analysis

We say that a pathway is mutated if there is at least one gene in the pathway carrying a somatic SNV or InDel. With respect to a particular pathway, each patient can be classified as CLL-/RS-, CLL-/RS+, CLL+/RS- or CLL+/RS+, depending on whether the patient is mutated in that pathway in none, at least one, or both of the CLL and RS phases, respectively. It follows that we can construct an *N* × 4 table **X** = {*x*_*ij*_} indicating the number of patients in each of the above groups in each among *N* pathways. We examined a total of 45 pathways in this study, which (after merging pathways provided by both KEGG and PanCancer) corresponds to *N* = 38. The total sum in each row of the table is obviously 17, the number of patients entering the study. The above table can be generated using a Multinomial-Dirichlet mixture model, as shown below:

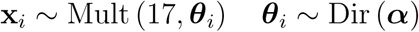

where **x**_*i*_ = (*x*_*i*1_, *x*_*i*2_, *x*_*i*3_, *x*_*i*4_) are the numbers of CLL-/RS-, CLL-/RS+, CLL+/RS- and CLL+/RS+ patients in pathway *i*, respectively, and ***θ***_*i*_ = (*θ*_*i*1_, *θ*_*i*2_, *θ*_*i*3_, *θ*_*i*4_) are the corresponding probabilities of observing each of the above four patient groups in pathway *i*. Given the posterior density of ***θ***_*i*_, we can estimate the posterior probabilities *P*_*i*1_ and *P*_*i*2_ that the RS and CLL phases harbour a SNV/InDel in pathway *i*, respectively, as well as the posterior probability *p*_*i*_ that RS is more likely than CLL to carry such an aberration, i.e. the probability that *P*_*i*1_ *> P*_*i*2_.

A similar analysis can be independently performed for the CNV data by assuming that a pathway harbours a copy number event, if there is at least one gene in the pathway overlapping with such an event. As above, we can calculate the posterior probability *q*_*i*_that the RS phase is more likely than the CLL phase to harbour a CN event in pathway *i*. The overall probability that RS is more likely than CLL to carry both a SNV/InDel and a CNV in pathway *i* is simply the product *p*_*i*_*q*_*i*_.

It follows that *π*_*i*_ = 1 − *p*_*i*_*q*_*i*_ is the probability of a Type I error, if we decide to call pathway *i* significantly more mutated in RS than in CLL. Subsequently, each pathway *k* in the set {*k*: *π*_*k*_ *π*_*i*_} is also called significantly more mutated in RS than in CLL, and the average Type I error rate (false discovery rate or FDR) over this set is equal to

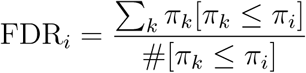

where [·] is the *Iverson bracket*. It is easy to see that the maximum possible value for FDR_*i*_ is *π*_*i*_, which is attained if *π*_*k*_ = *π*_*i*_ for all *k*. In other words, this probability also serves as an upper bound for the FDR, should we choose to label all pathways *k* satisfying *π*_*k*_ ≤ *π*_*i*_ as significantly over-mutated in RS (see also [4, 5] for an application of the same logic in differential expression analysis). In practice, we only consider a pathway *i* as significant, if *π*_*i*_ ≤ 5% (which corresponds to *p*_*i*_*q*_*i*_ 95%), in order to ensure an FDR over all discoveries not higher that 5%.

A final detail in the above model is the treatment of the vector of concentration parameters ***α***. We have examined three di***α***erent scenarios: (a) ***α*** = (1, 1, 1, 1), which corresponds to a flat Dirichlet distribution, (b) ***α*** = (*α*_1_, *α*_2_, *α*_3_, *α*_4_), with log *α*_*j*_ ∼ 1 for *j* ∈ {1, 2, 3, 4}, i.e. the logarithm of each element of the vector follows a uniform distribution in the interval (− ∞, + ∞), (c) a third option is to consider the mean and variance of each element 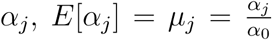 and 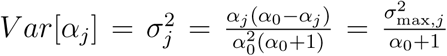, respectively, where *α*_0_ = Σ_*j*_ *α*_*j*_ and 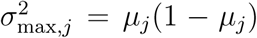. Instead of imposing a prior on *α*_*j*_ directly, we impose uniform priors on its mean and variance, *µ*_*j*_ ∼ *𝒰* (0, 1) and 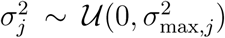 Among these three alternatives, it turns out that (a) is the most conservative, i.e. it identifies the smallest number of pathways as differentially mutated, and for this reason it is preferred.

### 2. Clonal analysis

We assume that each tumour sample (CLL or RS) is a mixture of normal (N) and cancer (C) cells, where the purity *ρ* of the sample indicates the proportion of cancer cells in the tumour. With respect to a particular somatic mutation *i*, there is a proportion *ϕ*_*i*_ of cancer cells (V) harbouring the mutation, while the remaining proportion of cancer cells (R) do not. The expected variant allele fraction or VAF, *f*_*i*_, for mutation *i* is given by the following formula:

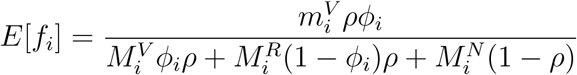

where 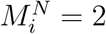 is the copy number of the normal cell population at the locus of mutation *i* (here, assumed to be 2), and 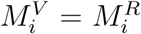 are the copy numbers of the R and V cancer cell populations at the same locus (here, assumed to be both the same). Finally, 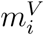 is the number of chromosomes in the cancer cells harbouring mutation *i*. This is assumed equal to 1, or equal to 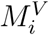, if the mutation is located in a loss-of-heterozygosity (LOH) region. Given *r*_*i*_ reads supporting the mutation (among a total of *R*_*i*_ reads covering the locus), we have:

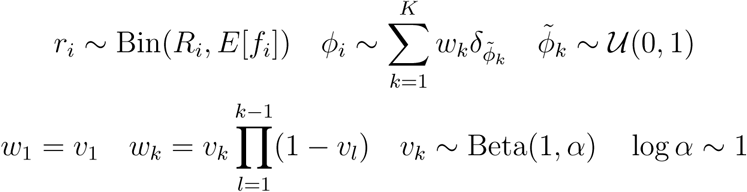

where 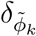 is a point-mass distribution centred at 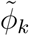. The above model assumes that the cancer cell fraction (CCF) *ϕ*_*i*_ for mutation *i* follows a discrete mixture distribution over 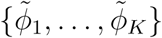, where *K* is a large number. The mixture weights *w*_*k*_ are generated from a stick-breaking process with concentration parameter *α*. The above model implies that all mutations in a particular sample share a small number of distinct CCF values, as expected in a tumour composed of homogeneous cell populations (i.e. *clones*).

In the above model, we calculated the posterior distribution of each CCF value *ϕ*_*i*_ in each sample using ADVI as implemented in `Stan`. Subsequently, we calculated the difference in the CCF values between each pair of CLL and RS samples for each somatic mutation. A difference between the matching boundaries of the 95% credible intervals of the CCF values larger than 0.25 from CLL to RS indicates clonal expansion, a difference of the same magnitude in the opposite direction indicates clonal contraction, while a difference of smaller magnitude in any direction indicates neither expansion nor contraction. Given this classification of mutations, we constructed an *N* × 3 table, where rows correspond to pathways and columns correspond to the above three mutation groups (i.e. expansion, contraction, neither). The elements of the table indicate the number of mutations in each group in each pathway. The sum across each row equals the total number of mutations in the corresponding pathway across all samples. This table was modelled using a Multinomial-Dirichlet mixture as in the previous section, and the posterior probabilities of clonal expansion and contraction were calculated for each pathway. Pathways for which clonal expansion was more likely than contraction with a probability higher than 95% (which corresponds to an FDR less than 5%; see previous section) were tagged as significant.

### 3. Enrichment analysis

As mentioned in the Overview section, genes with FDR less than 1% were considered differentially expressed. With respect to a particular pathway, a gene can be classified as differentially expressed and in the pathway, differentially expressed but not in the pathway, not differentially expressed and in the pathway, and not differentially expressed and not in the pathway. As in the two previous sections, we constructed an *N* × 4 table, where rows correspond to pathways and columns to the above gene categories. Elements of the table indicate the number of genes in each pathway in each category, and the total sum across each row is equal to the number of genes that entered the differential expression analysis. As in the previous sections, we employed a Multinomial-Dirichlet mixture model, and we calculated posterior probabilities for each gene category in each pathway. If the proportion of differentially expressed genes in the pathway was higher than the proportion of differentially expressed genes not in the pathway with probability higher than 95% (FDR less than 5%), then the pathway was tagged as enriched in differentially expressed genes.

